# InfoDengue: a nowcasting system for the surveillance of dengue fever transmission

**DOI:** 10.1101/046193

**Authors:** Cláudia T Codeço, Oswaldo G Cruz, Thais I Riback, Carolin M Degener, Marcelo F Gomes, Daniel Villela, Leonardo Bastos, Sabrina Camargo, Valeria Saraceni, Maria Cristina F Lemos, Flavio C Coelho

## Abstract

This study describes the development of an integrated dengue alert system (InfoDengue), operating initially in the city of Rio de Janeiro, Brazil. It is a project developed as a partnership between academia and the municipal health secretariat. At the beginning of each epidemiological week, the system captures climate time series, dengue case reporting and activity on a social network. After data pre-processing, including a probabilistic correction of case notification delay, and calculation of dengue's effective reproductive number, indicators of dengue transmission are coded into four dengue situation levels, for each of the city's ten health districts. A risk map is generated to inform the public about the week's level of attention and the evolution of the disease incidence and suggest actions. A report is also sent automatically to the municipality's situation room, containing a detailed presentation of the data and alert levels by health district. The preliminary analysis of InfoDengue in Rio de Janeiro, using historical series from 2011 to 2014 and prospective data from January to December 2015, indicates good degree of confidence and accuracy. The successful experience in the city of Rio de Janeiro is a motivating argument for the expansion of InfoDengue to other cities. After a year in production, InfoDengue has become a unique source of carefully curated data for epidemiological studies, combining epidemological and environmental variables in unprecedented spatial and temporal resolutions.

**Ethical committee approval:** 26910214.7.0000.5240

## Introduction

Dengue fever transmission is characterized by significant inter-year variability with seasons of intense activity separated by periods of very low to no detectable activity. Complex interactions between environmental factors (such as temperature and humidity), human factors (such as population immunity and mobility) and viral factors (circulating strains) modulate the transmission of dengue. This complexity leads to pronounced prediction uncertainties making it hard to prepare for and allocate resources to reduce disease burden.

Currently, there is a global effort to improve the sensibility and speed of disease surveillance systems by various means (L’Azou et al. 2014): by developing multivariate methods which bring together information from different sources; by incorporating alternative sources of information such as symptom report in social networks (Milinovich et al. 2014), or the monitoring of search terms in search engines (Chan et al. 2011) and by adopting variables not directly associated with the transmission such as meteorological variables (Coelho and Carvalho 2015).

Examples of new surveillance approaches for dengue are found in Singapore, Philippines and Cambodia (Huy et al, 2010). In Singapore, a web-based alert system (www.dengue.gov.sg) classifies sites in terms of transmission risk: low, medium or high. Risk is determined by the presence of clusters of cases, defined by two or more cases of dengue occurring within 14 days within the same locality. An alert map with the case clusters is made available to the population to trigger actions against dengue. In 2013, the government of the Philippines launched an online system (www.dost.gov.ph) through which the population can check the risk of dengue at each locality based on weekly monitoring of mosquito populations, carried out by 45 thousand public schools throughout the country. Before the school term starts, the government distributes egg traps with larvicide to all schools. Each week, the school coordinator reports how many traps are positive, and this amount is translated into colored flags. In most cases, dengue surveillance systems focus on gathering direct evidence of transmission for situational awareness and/or informing control strategies.

Rio de Janeiro is a tropical city with ca. 6.5 million inhabitants within a metropolitan region with ca. 12.1 million inhabitants (IBGE, 2014); the hottest and humid season comprehend the period from November to April, and the colder and drier from May to October (Câmara et al, 2009). Dengue fever is endemic in Rio de Janeiro since 1986-1987, when DENV-1 arrived and caused high disease burden, with more than 1 million reported cases. The first isolation of DENV-2 occurred in 1990, accompanied with the first cases of severe dengue; after this period was responsible for an outbreak between 2007 and 2008 (Teixeira et al. 2009, Fares et al. 2015).

The occurrence of DENV-3 was first reported in 2000, and in 2002 it was responsible for a large epidemic with more than 280.000 reported cases (Nogueira et al. 2001, Fares et al. 2015). The presence of DENV-4 was detected in 2010 (Nogueira and Eppinghaus 2011, Fares et al. 2015) and currently, DENV-1 and DENV-4 are the most prevalent serotypes circulating in Rio de Janeiro (Fares et al. 2015). Due to economic and touristic importance, the city receives a large daily influx of people from different regions, a situation that may increase the risk of entry and dissemination of new diseases (IBGE 2010, Nogueira et al. 2006, Nogueira and Eppinghaus 2011). High heterogeneity and urban complexity makes surveillance and control of vector-borne diseases an immense challenge.

Dengue surveillance and control activities are informed by periodic larval surveys (3-4 per year) that are used to rank areas according to Aedes aegypti infestation levels; and control charts are used to identify excess of notified cases. In Rio de Janeiro, these data are analyzed weekly in the city's Dengue Situation Room. The aim of this paper is to describe the implementation and first year of operation of a new method, the InfoDengue nowcasting system, used to improve the continuous monitoring of dengue fever in Rio de Janeiro, at a useful scale for health management. Integrating readily available data from different sources, types and spatiotemporal resolution, this system was implemented and is operational in the city of Rio de Janeiro, Brazil, since January 2015, providing a public website (info.dengue.mat.br) with the status of the dengue incidence, which is weekly updated, and a detailed report for the city's dengue situation room.

The key concept behind InfoDengue is “transmission”, measured in terms of the effective reproductive number (Rt). In theory, Rt is measured as the mean number of secondary cases generated by a primary case at a time t. A number greater than one implies sustained transmission, which is important information for public health decision. Our transmission-based surveillance system has four levels, coded in a green-yellow-orange-red color scale (Table 1). In the following sections, we present the development of the system, followed by a description of its operation during its first year in Rio de Janeiro.

**Table 1.**
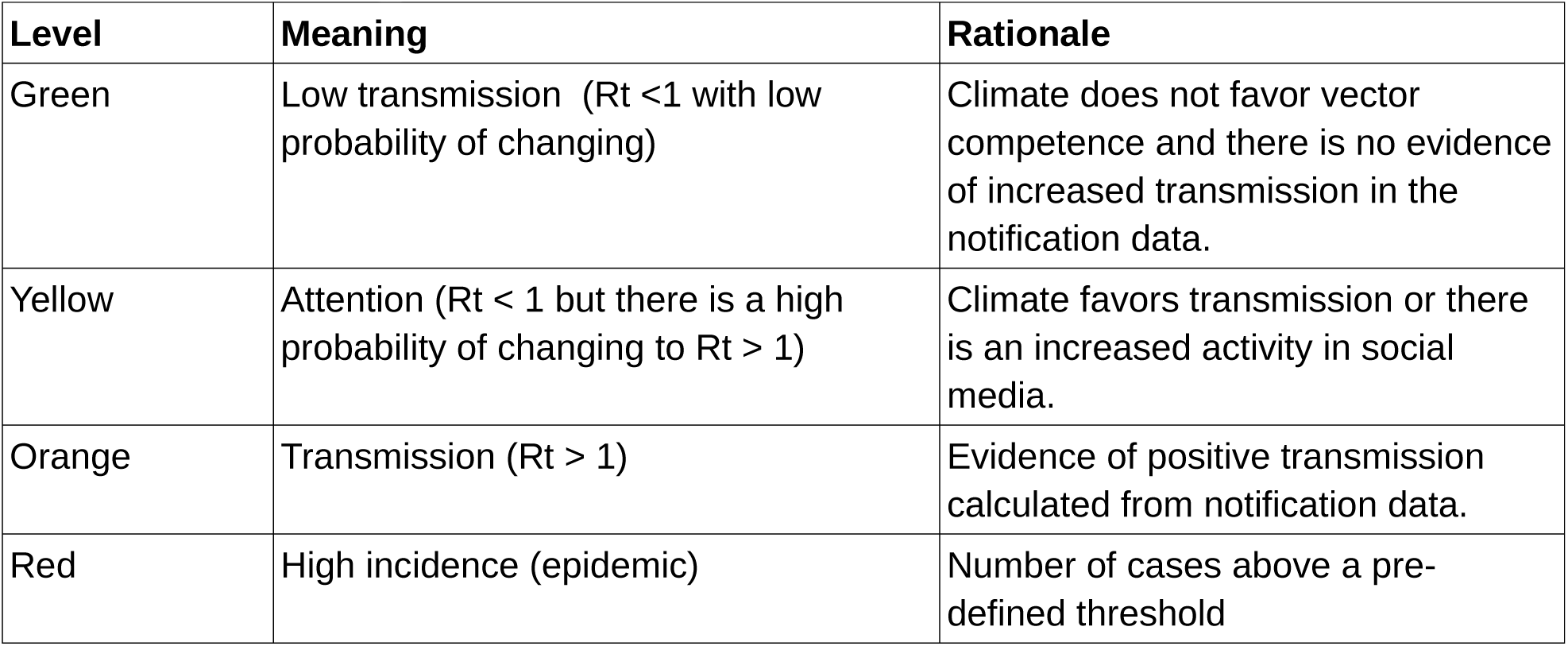
Levels of the InfoDengue system

## Methods

### Study site

Rio de Janeiro city (22.9068 S, 43.1729 W) has a population of 6.5 million inhabitants distributed in an area of 1200 km2. Due to its size, dengue control and monitoring activities are structured in 10 health districts (Áreas Programática da Saúde) (Figure 1 and Table 3). AP1 is the downtown area, AP2.1 and AP4 are located at the seashore and house a population with average to high income; AP2.2, AP3.1, AP3.2, AP3.3 are in the northern region, and are a mixture of very poor and middle class neighborhoods; AP5.1, AP5.2 and AP5.3 are located in the periphery, mostly poorer neighborhoods that are strongly connected to the neighboring cities of the Rio de Janeiro metropolitan region. The 10 Health Districts also have distinct climates, depending on their position in relation to the sea, bay, and mountains that cross the city.

**Figure 1.**
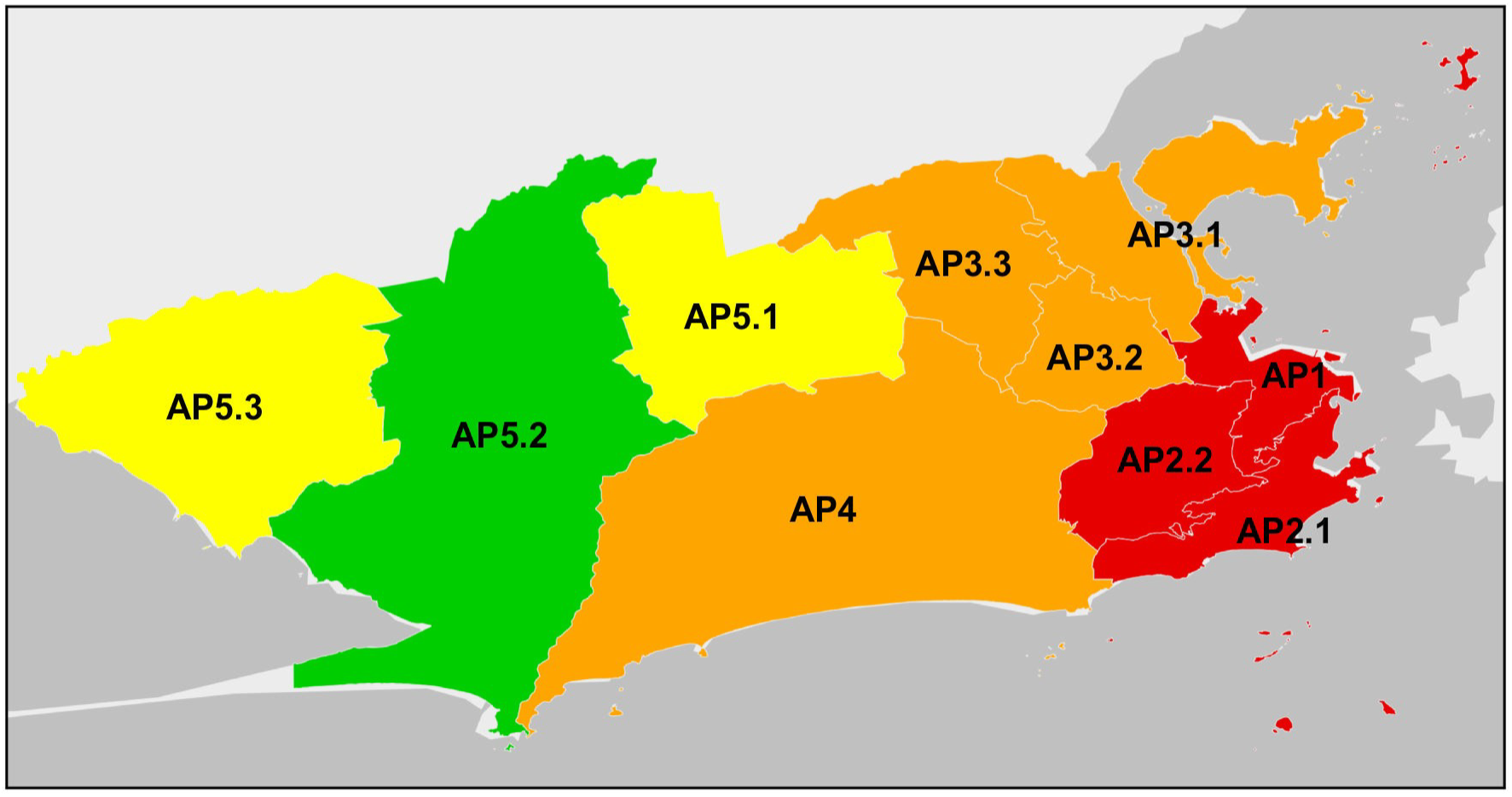
Division of Rio de Janeiro city into its ten health districts. See table 1 for further description.

### Data

A dataset containing time series of air temperature, dengue notifications, and tweets on dengue from January 2010 to December 2014 in Rio de Janeiro was used to derive a set of rules for the alert system. Climate data consisted of minimum weekly air temperature gathered from 4 meteorological stations located at the airports (Table 2). Messages on twiter indicative of having dengue and georeferenced to Rio de Janeiro were provided by the Observatorio da Dengue at the Federal University of Minas Gerais (UFMG) who carries out automatic message classification to remove messages mentioning dengue in other contexts, as described elsewhere (Gomide et al. 2011). Reported suspected cases of dengue were obtained from the Brazilian National Notification System (SINAN and DENGON). The following variables were obtained: date of symptom onset, date of notification, date of database entry, and neighborhood of residence within Rio de Janeiro. Notification data were aggregated by the 10 health districts.

**Table 2.** Rio de Janeiro city is divided into 10 health districts. This table shows the population size, dengue's 5 year attack rate, % of weeks with Rt > 1 and the meteorological station associated to each area (Airport codes).

| Health district | Population | 2010-2014 dengue attack rate (x100) | % weeks with $R_t > 1$ (2010-2014) | Meteorological station |
| --- | --- | --- | --- | --- |
| AP1 | 226,963 | 4.95 | 14.7 | SBRJ |
| AP2.1 | 552,691 | 4.81 | 20.1 | SBRJ |
| AP2.2 | 371,120 | 3.34 | 17.1 | SBRJ |
| AP3.1 | 735,788 | 2.43 | 18.2 | SBGL |
| AP3.2 | 489,716 | 3.90 | 18.6 | SBGL |
| AP3.3 | 924,364 | 4.17 | 17.8 | SBGL |
| AP4 | 838,857 | 2.85 | 17.0 | SBJR |
| AP5.1 | 655,874 | 6.56 | 20.9 | SBAF |
| AP5.2 | 665,198 | 4.24 | 19.7 | SBAF |
| AP5.3 | 368,534 | 3.18 | 16.3 | SBAF |
| **Whole city** | **5,829,105** | **3.99** | **18** |  |

### Correction of the delay in case notification

Before proceeding with the analysis, dengue notification delay had to be fixed. **Typically the SINAN database remains open for six months to update case counts retrospectively. Delays reflect the time taken for a patient to visit the doctor, the time the doctor takes to fill in the notification form, and the time taken for a technician to type and upload the form to SINAN. We developed a probabilistic model to estimate the number of cases at time t from incomplete case reports, considering that information at time t is partial (censured) and only will become available in the future. In other words, we want to predict the number of cases at time t that will be known for certain only 6 months ahead. The probabilistic model is detailed in the Appendix**. Figure 2 shows the agreement between estimated and true case numbers using this model.

**Figure 2.**
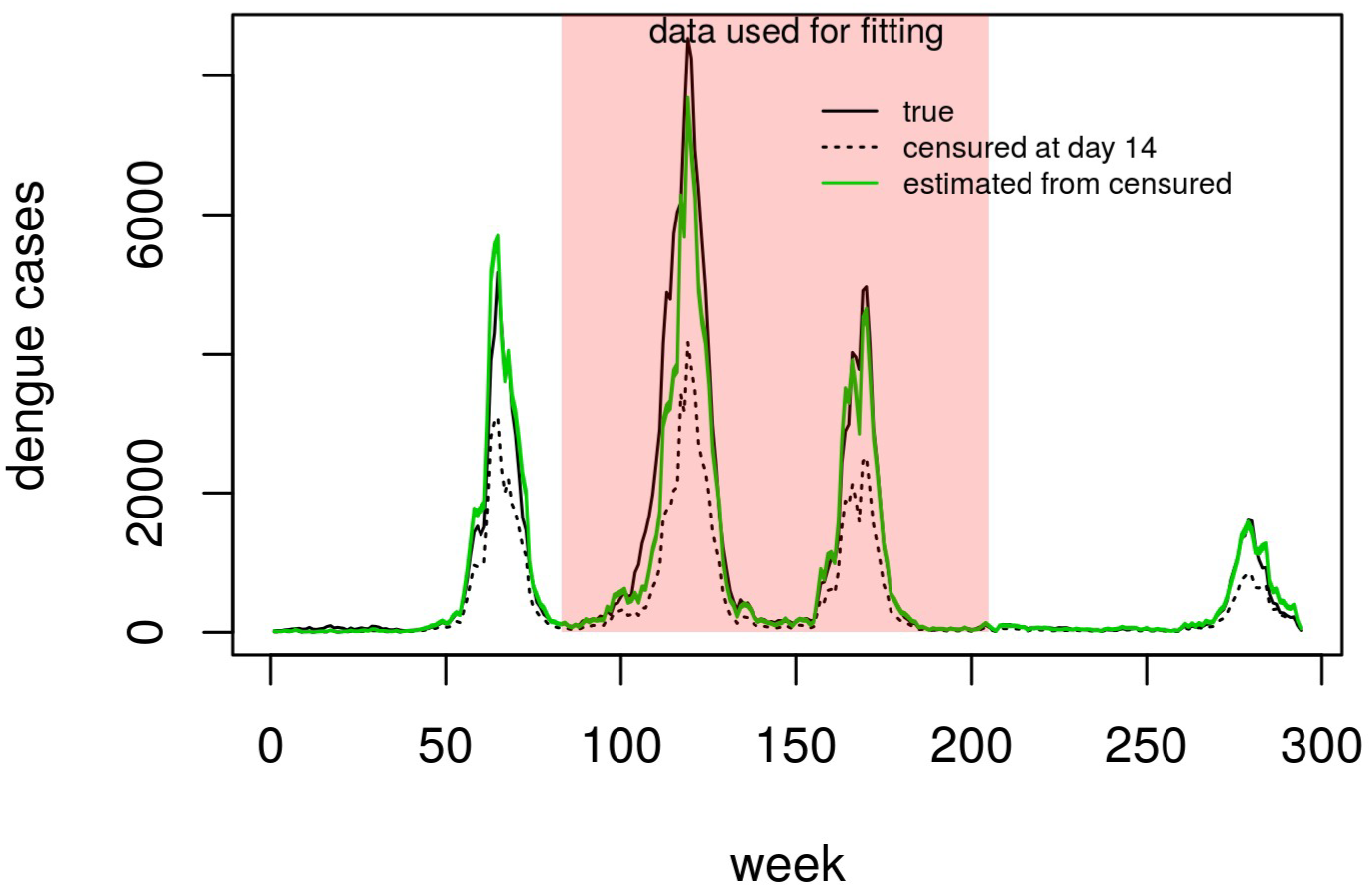
Goodness-of-fit of the probabilistic model used for correcting the notification data for the delay between disease onset and entry in the database. The dashed line is the fraction of cases that are notified within 2 weeks from its occurrence. The green line is the number of cases as estimated by the model. The black line is the total number of cases that were notified for that week (only known 6 months later). The model was fitted to the data in the shaded area and validated in the subsequent time window.

### Measuring disease transmission

As said before, the core concept in InfoDengue is “transmission”. In other words, we want to identify periods of critical (Rt > 1) and subcritical transmission (Rt < 1). To estimate Rt from incidence data, after correcting for delay, we employed Wallinga and Lipsitch (2007)'s equation:

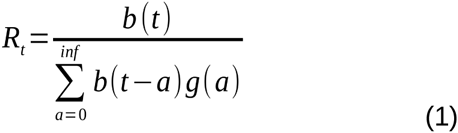

where b(t) is the corrected case count at week t, and g(a) is the distribution of dengue's generation interval (defined as the time between symptoms onset in a primary case and symptoms onset in a secondary case). For simplicity, we assumed that g(a) follows a delta distribution with mean of 3 weeks, the underlying assumption being that all secondary infections of a primary case occurred at an interval exactly equal to the mean generation interval (Wallinga and Lipsitch 2007). Three weeks is approximately the sum of the average intrinsic and extrinsic incubation periods of dengue at temperature 25C (6 + 13 days, respectively). With g(a) being a delta distribution, equation 1 is equivalent to the Stallygrass estimator, and credible intervals for Rt can be computed using the method described in Coelho and Carvalho (2015). For declaring Rt > 1, we considered a cutoff of p(Rt > 1) = 0.9.

Figure 3 shows the time series of notified dengue cases in each of the 10 health districts of Rio de Janeiro, from January 2010 to December 2014, marking the weeks with Rt > 1 (grey vertical bars). We observed Rt > 1 in ca. 17-20% of the weeks, mostly concentrated in the period between February and May (late summer - early fall). Isolated week estimates of Rt are quite volatile. To avoid raising false alarms, an orange alert indicating sustained transmission was only issued after 3 consecutive weeks with Rt > 1. This period corresponds to one generation time which is the natural scale for dengue dynamics.

**Figure 3.**
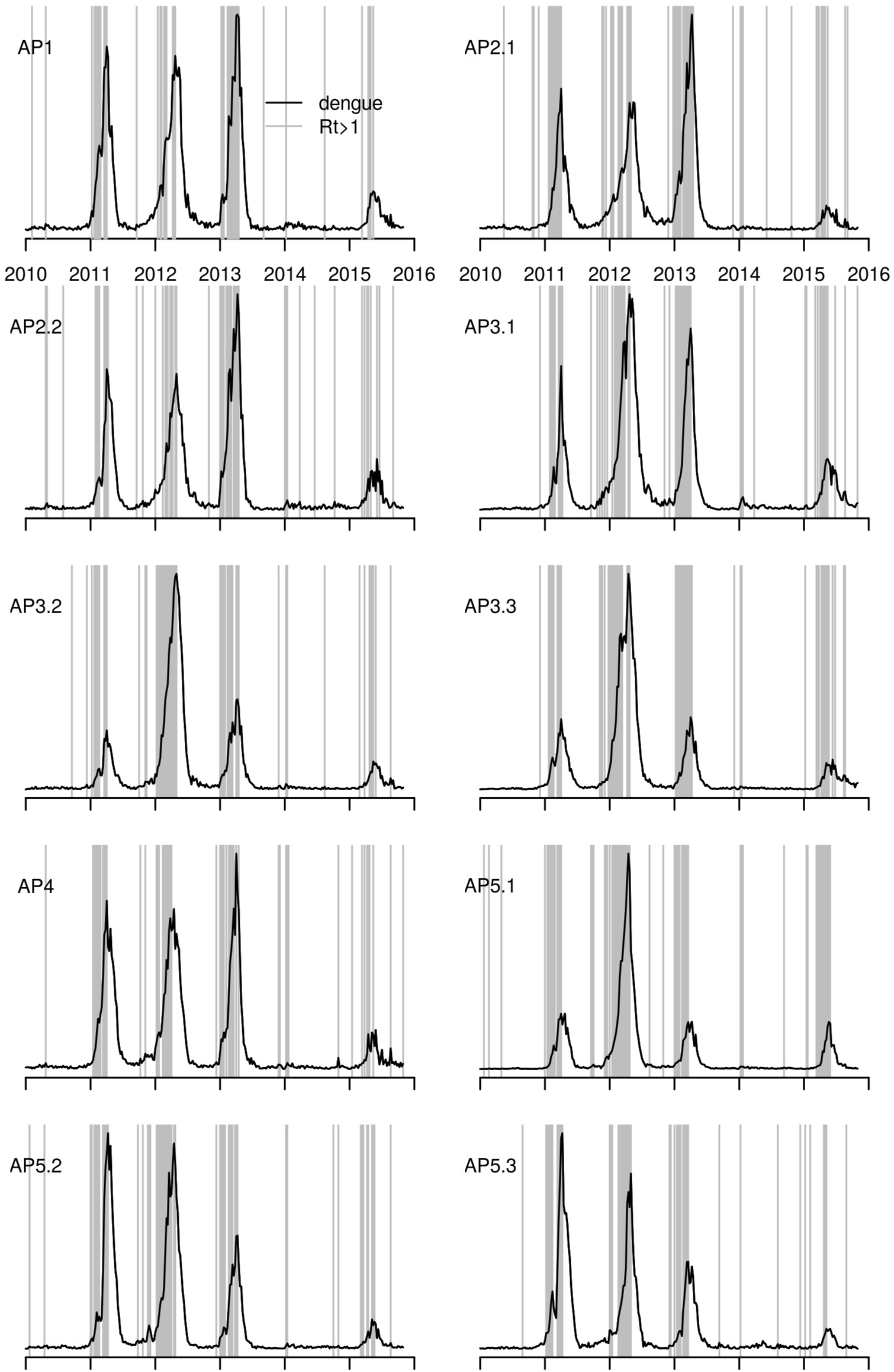
Time series of dengue notification in the 10 health districts of Rio de Janeiro (Jan 2010 – Dec 2014). The grey lines indicate weeks with Rt > 1 (p-value < 0.1).

### Reproduction rate × temperature

To study the association between temperature and dengue transmission, the city was divided into 4 sub-areas (corresponding to the health districts under the influence of each meteorological station, as in Table 2). In each of the four sub-areas, Rt was calculated from local incidence data as described above. Figure 4A compares the distribution of temperature in weeks with critical and subcritical transmission. The boxplots are similar among health districts 1 to 4, suggesting a single common temperature cutoff to discriminate critical and subcritical weeks. To identify this cutoff, a ROC curve was fitted to each of the four temperature-dengue datasets (Figure 4B). A cutoff point at 22C presented sensitivity above 80% to detect Rt > 1, with reasonable specificity in the health districts 1 to 4.

**Figure 4.**
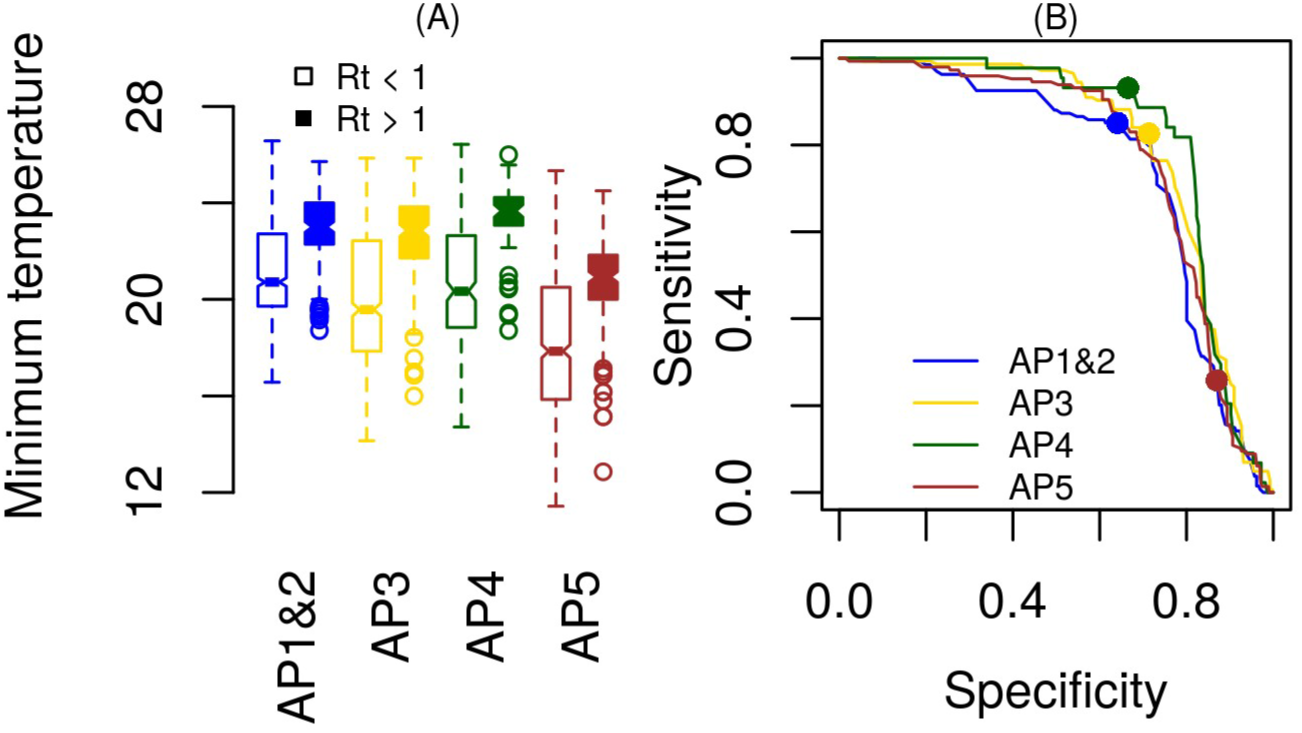
(A) boxplots of temperature values in weeks with Rt above or below 1. Each color corresponds to a different area of the city. (B) sensitivity-specificity plot of different cutoff points of temperature to discriminate weeks with Rt > 1. Dots indicate the cutoff used (22C).

For Health Districts 5.x, a 22C cutoff is too high. This district also had significantly lower temperatures than the other areas. No geographical feature of the area explains this difference in temperature, and we wonder if this could be due to some specificity of the meteorological station of the airport. For simplicity, we kept the same cutoff point for all districts, and the effect of this decision is discussed later.

### Tweeting is linearly associated with dengue incidence

Twitter is a realtime source of information on dengue symptoms activity in a population. Tweeting on dengue showed strong correlation with the number of notified cases (Figure 5A. Pearson's r = 0.75, p < 0.001). Looking at the time series, however, it is clear that the association is stronger during the increasing and decreasing phases, than during the disease peaks (Figure 5C), emaning that epidemic peaks are not correctly captured by the tweets.

**Figure 5.**
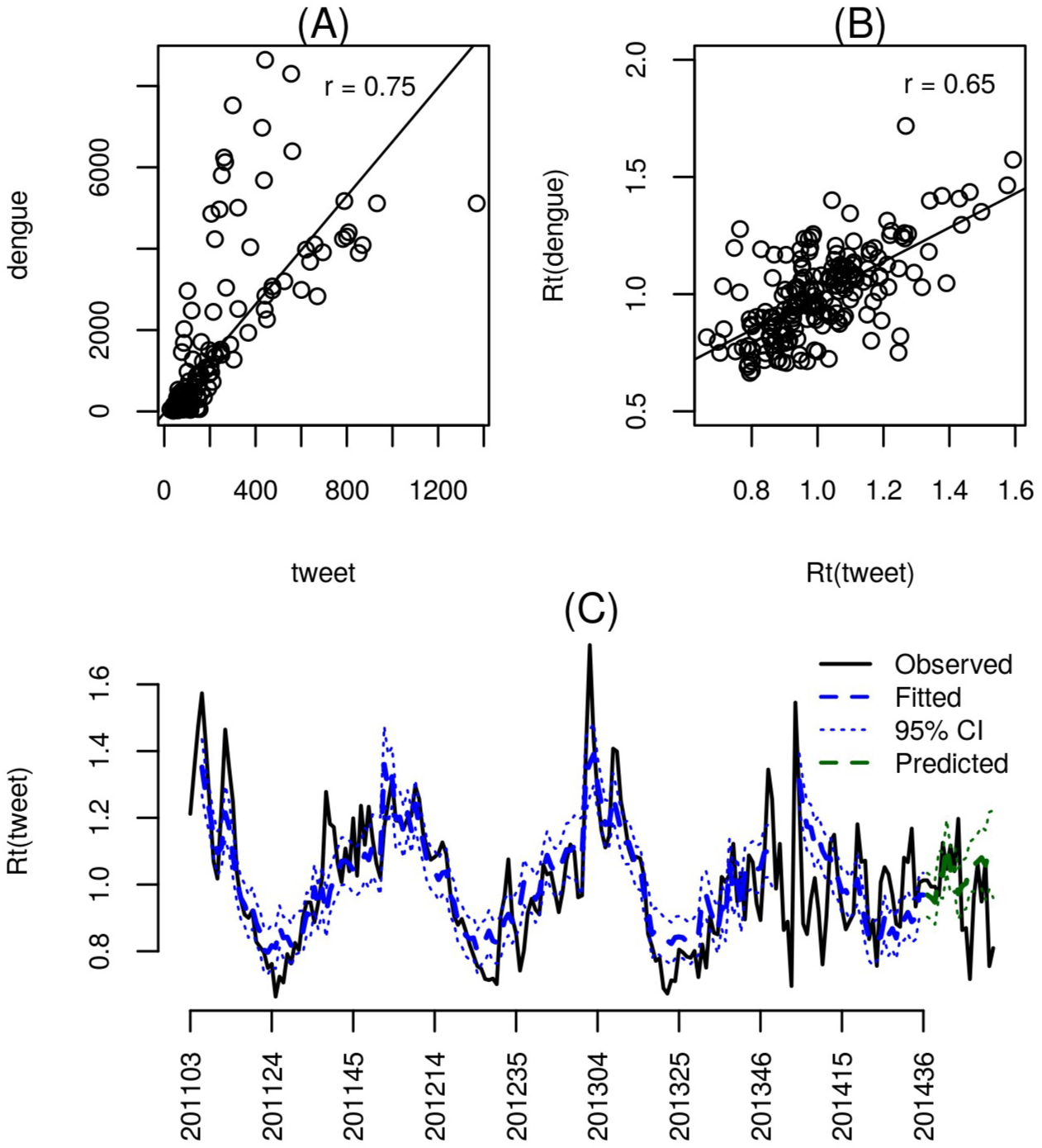
Time series of dengue notifications in the city of Rio de Janeiro (Jan 2010 – Dec 2014) and the number of twits indicative of dengue symptoms during the same period.

As an alternative, we considered the computation of Rt(tweet) calculated as if tweets were the actual cases of disease, using equation (1). The Pearson's correlation between Rt(dengue) and Rt(tweet) is somewhat smaller (Figure 5B, Pearson's r = 0.65, p < 0.001), but the relationship is more linear. We therefore investigated the association between Rt(dengue) and Rt(tweets), by fitting regression models. A gaussian additive mixed model was required to proper fit the relationship between the reproductive numbers of dengue cases and tweets.

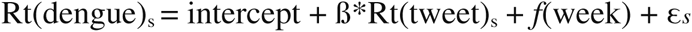

An auto-regressive term of order 1 (AR-1), which models the residual at time s as a function of the residual of time s-1 and noise (εs = ρεs–1 + qs) was included in order to account for the significant autocorrelation, that was present in a previously adjusted model without autoregressive term (Phi=0.60). The final model has β = 0.27 (SE=0.054, p<0.001).

In the alert system, the Twitter time series is in the following way: a significant increase in social media activity (measured as Rt(tweet) > 1) is used as a warning (yellow alert). More sporadically, when the notification dataset is offline, the number of tweets is used to infer the number of cases using a linear regression model fitted to the last one year of data.

## The InfoDengue pipeline

The analysis described above suggested a strong association between temperature, twits and dengue and the feasibility of developing a nowcasting system for dengue transmission using these data. An analytic pipeline was developed and implemented as shown in Figure 6.

**Figure 6.**
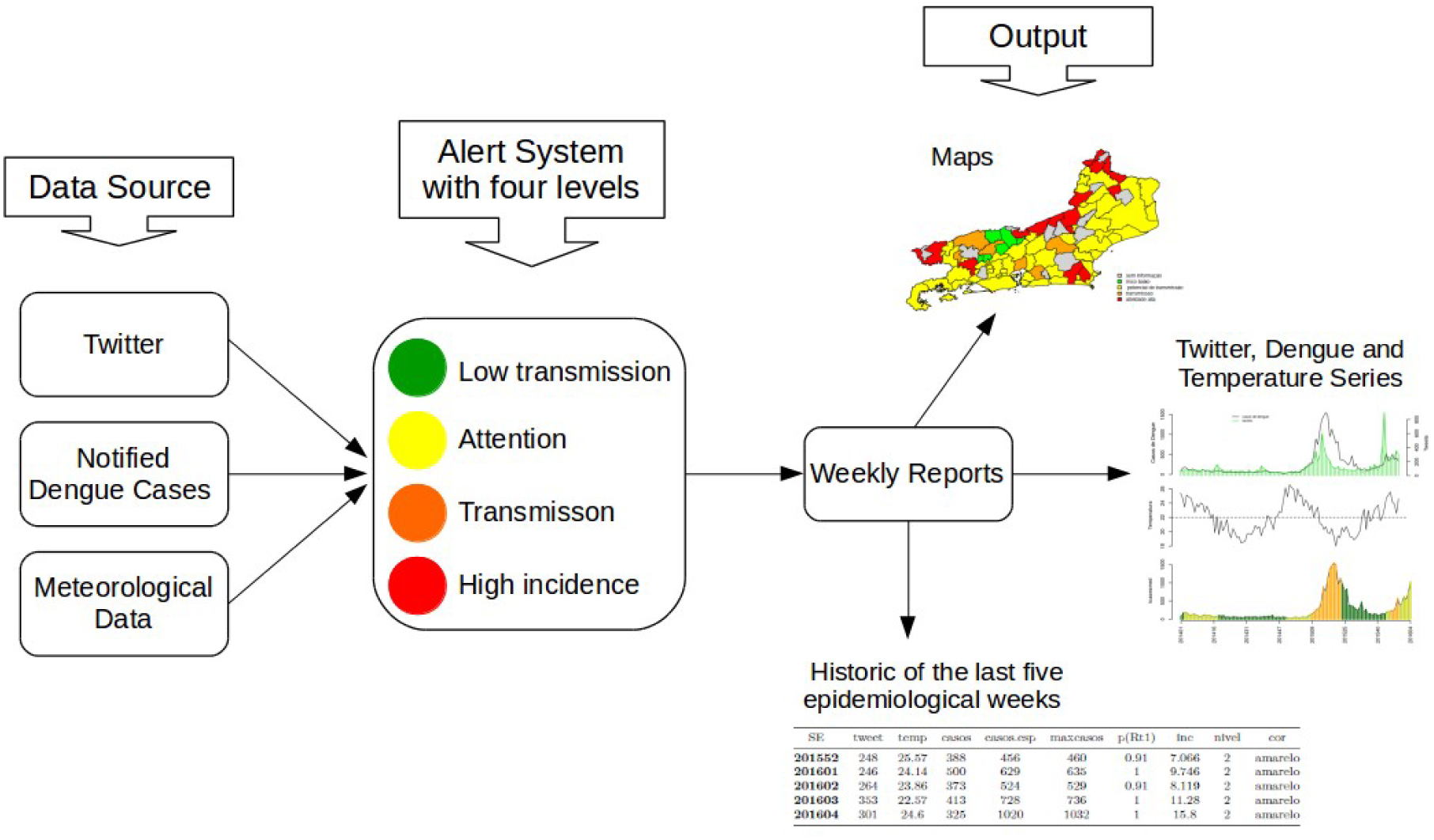
InfoDengue pipeline

At the beginning of each new week, the pipeline receives an updated value of minimum temperature (Tmin), number of tweets (Tw) and estimated number of cases (*Y*), per health district. Based on these data, a set of rules is applied to define the alert level.

A = 1 if Tmin > 22 for 3 consecutive weeks, 0 if otherwise
B = 1 if Rt(tweet) > 1, with probability > 0.9 for 3 consecutive weeks, 0 if otherwise
C = 1 if Rt > 1 with probability > 0.9 for 3 consecutive weeks, 0 if otherwise
D = 1 if estimated incidence > 100 cases per 100.0000 inhabitants, 0 if otherwise

with these rules, we build the color code system (Table 3).

**Table 3.** Confusion matrix showing the agreement between the classification of dengue risk proposed by specialists and by the automated rule system.

| Classification |  | InfoDengue |  |  |  |
| --- | --- | --- | --- | --- | --- |
|  |  | Green | Yellow | Orange | Red |
| Specialist | Green | 0.807 | 0.082 | 0.060 | 0.049 |
|  | Yellow | 0.567 | 0.290 | 0.142 | 0 |
|  | Orange | 0.058 | 0.058 | 0.783 | 0.003 |
|  | Red | 0 | 0.011 | 0.011 | 0.977 |

### Confusion matrix

To measure the adherence of the proposed rules to a gold standard, we asked two specialists to manually classify the incidence series from 2011 to 2014, according to our 4-level alert system. The same period was also classified using the automated methodology. The result is presented in the form of a confusion matrix, *C*, whose elements *c*_*ij*_, are the fraction of weeks classified by the specialist as *i* and by the system as *j*. So in a perfect system we would have the main diagonal of the matrix composed of just ones while the remaining elements are zero.

### First year of operation

The system was launched in January 2015. To assess the performance of the system between weeks 201501 and 201544, we first analyzed the quality of the notification delay correction. Dengue data with and without correction (for the delay) were compared using the following measurement of error:

Without correction: error(w) = (all reported cases with onset at week w – reported cases with onset at week w, known at week w+1)

With correction: error(w) = (all reported cases with onset at week w – estimated cases using correction model)

Secondly, the alert level provided at real time was compared to the level ascertained retrospectively, after complete information was collected. This comparison is only qualitative, since the time series is still short for a more formal statistical analysis.

## Results

Figure 7 shows the time series of dengue cases for each Health District, from Jan 2011 to Dec 2015 (note that the system was prospectively operated from Jan 2015 on). The colors indicate the level of alert defined by the InfoDengue rule system. In general, the system moved gradually from green to yellow to orange and, in some cases, to red. This is the desirable state of a warning system. In the Health Districts 5.x, mainly in 2012, the triggering of the orange level was not preceded by the yellow alert. This suggests that the temperature cutoff was actually to high for this area, as already predicted by the ROC analysis (Figure 4). In 2014 the system went only up to yellow alert, indicating adequate climatic conditions for transmission and lack of an actual incidence increase. This was one of the driest years in the last decade, a possible explanation for the unusually low dengue transmission.

**Figure 7.**
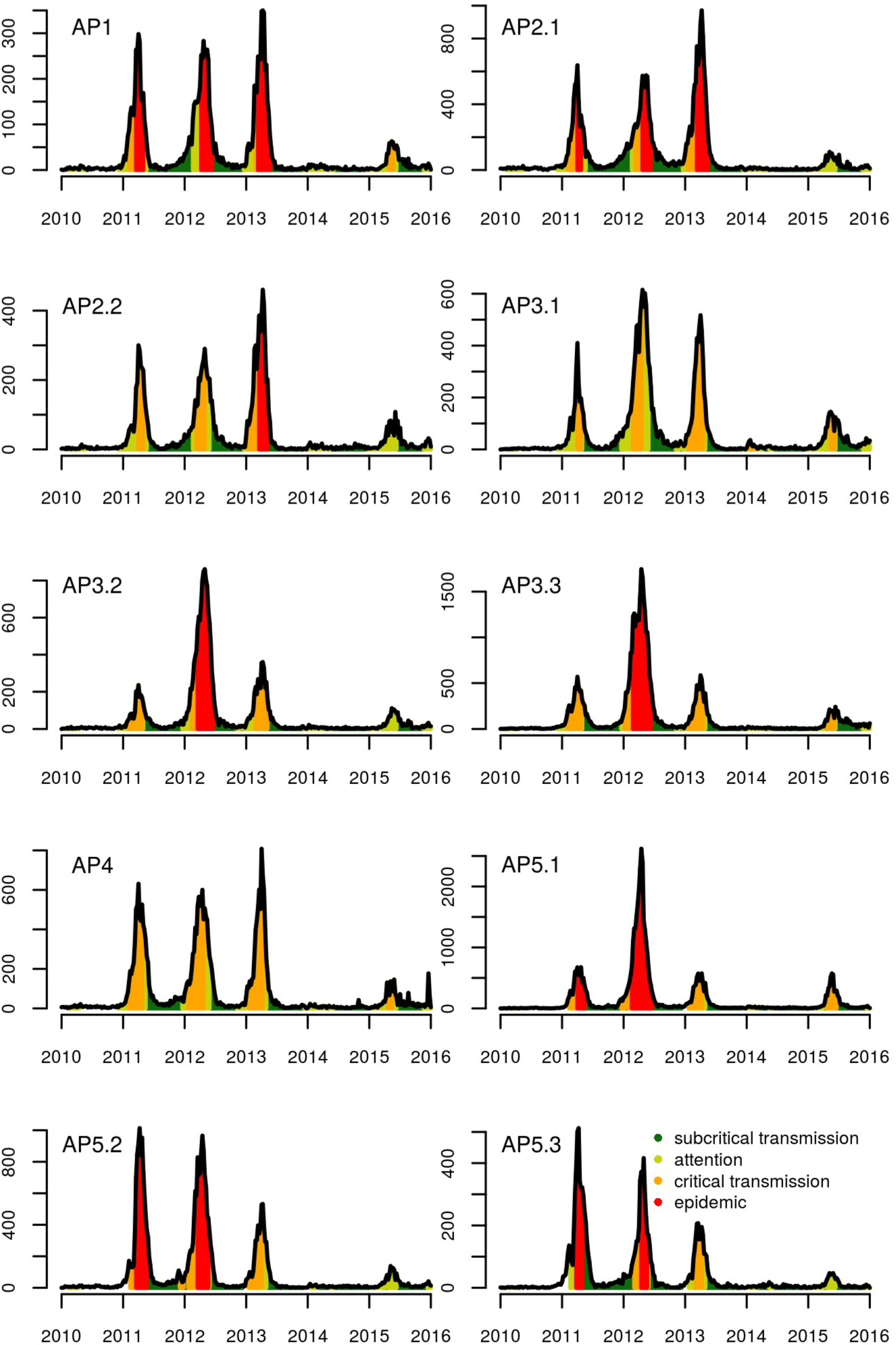
Time series for notified cases of dengue in Rio de Janeiro's health district APS 1 and the classification of the alert levels generated by InfoDengue.

In general, the automated nowcasting system displayed good adherence to the post-hoc classification by specialists, with agreements of 87% in green, 78% in orange and 97% in red weeks (Table 3). The yellow level had the lowest agreement, 29%. More specifically, InfoDengue classified as Green, 57% of weeks that a specialist would classify as Yellow. This means that the system was less sensitive than the specialist classification. Since the yellow level is the first wake-up call for health care workers, in principle the more sensitive the better. On the other hand, a system that overemphasizes sensitivity, at the cost of reducing specificity, might loose credibility. It is important to note that, in deciding the color level, specialists had access to the full case report time series. This means that, while deciding the level at a given week, they had information with respect to future weeks. Since our alert system is used for nowcasting, it only has historical and current data to base its decision on. For that level, this poses a particular challenge to rely only on reported cases, which lead us to adopt complementary environmental data.

With respect to reported cases, if there is enough sustained transmission the system will already issue at least an orange alert. What triggers the yellow level is when that situation is not yet present but environmental conditions are prone to its occurrence ‐‐ be it favorable climate for mosquito activity, be it significant attention level on social media ‐‐, factors that were not taken into account by the specialists. The later is considered since significant activity in social media combined with low case report can indicate higher underreporting. Nonetheless, we are working on enhancements on that particular level for better agreement. A possible alternative would be to incorporate forecasting into the model, which is a challenge in itself.

### Dengue seasonality

Figure 8 shows the seasonality of dengue transmission in Rio de Janeiro, according to our models. The dengue season (orange + red) is well contained within the warm season indicated by the yellow area. Sustained transmission tends to occur from late January to late April, and the epidemic season is concentrated between March and May.

**Figure 8.**
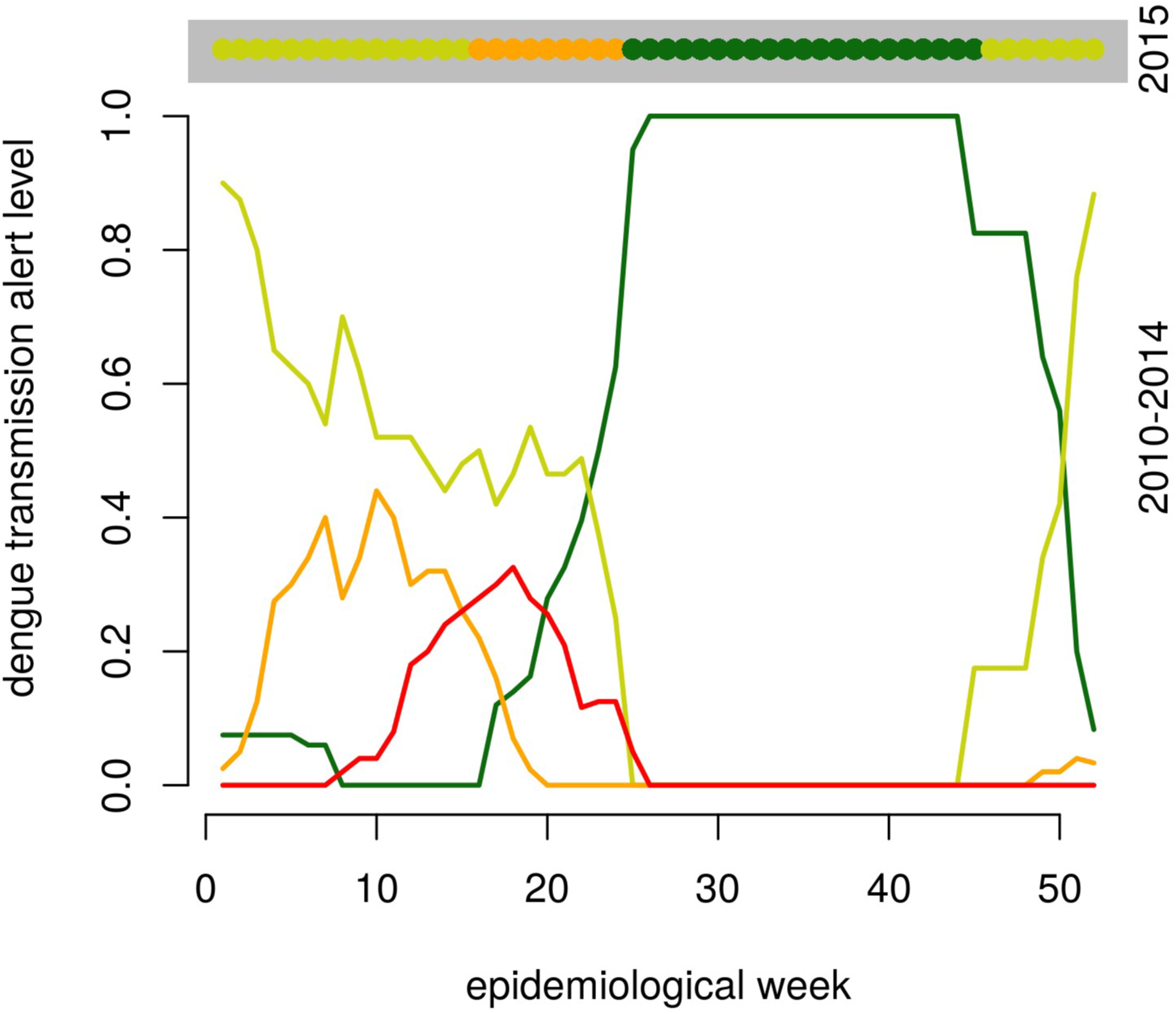
Seasonality of dengue transmission in Rio de Janeiro, from 2010 to 2014. Green means low transmission risk; yellow means proper conditions for dengue transmission; orange means evidence of sustained transmission; red means high dengue activity (above 100 cases:100,000 inhabitants). The top figure, within the grey area, shows the 2015 transmission pattern of suspected dengue. Later, we came to know that an unknown fraction of these cases were actually Zika virus infections.

### Assessment of the first year of operation

A total of 20,773 suspected cases of dengue were reported in 2015. Due to reporting delay, only 23.8% of the cases were known in the first week from occurrence. The error introduced by this delay is seen in Figure 9 (left boxplot), which shows the distribution of the difference between known-cases minus all yet-to-be-known cases. This difference is mostly negative, but can be positive as sometimes some suspected cases are discarded (infrequent). The delay correction procedure provided an unbiased estimation of the yet-to-be-known cases (Figure 9 right boxplot). The average error was of -3 cases, in comparison with -29 for the uncorrected estimator. Contrasting with the crude measurement, the estimated incidence both overestimated and underestimated the number of cases. In practice, both corrected and uncorrected measurements of incidence were included in the weekly reports.

**Figure 9.**
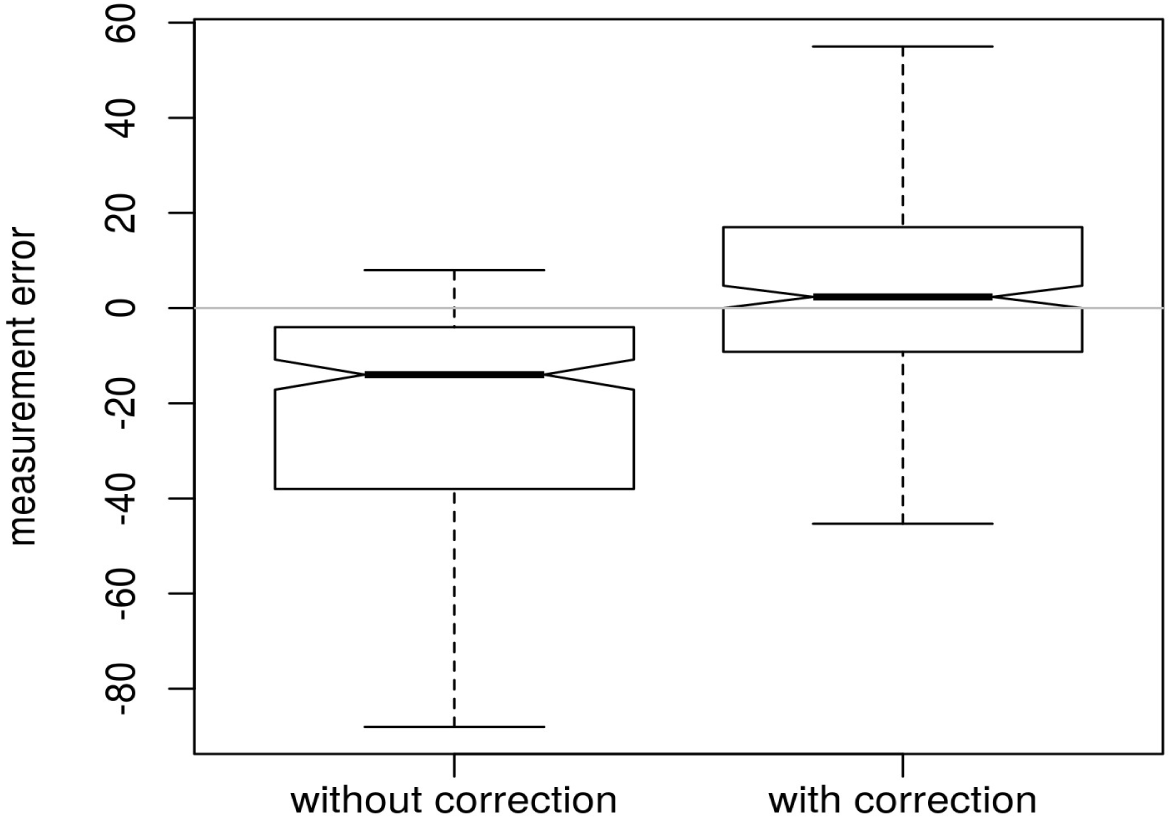
Performance of InfoDengue during the 2015 season. Left boxplot: distribution of the cases missed by the reporting delay as measured by the difference between all cases eventually reported, and those reported readily in the first week. Right box: distribution of the same measurement error, after applying the delay correction model (see text for details).

During the first year of operation, Rio de Janeiro was in yellow alert from January to mid March, due to the summer temperatures. During the first two months 30-60 cases/week occurred, but in March, incidence started to increase steadily reaching 1000-1500 cases/week between April-June. Sustained transmission (orange alert) was ascertained for the first time in April 29 and remained so until June 24. Only the Health Districts APS 3.1 and 3.3 stayed Orange during the whole period, the remaining shifted between Orange and Yellow. APS 1, 2.1 and 5.3 were the least affected (only 0 - 2 weeks with orange flag). After June 24, all Health Districts return to the Green state, with a stable incidence level of 200-300 cases/week during the winter and spring months of 2015. No red alert was raised during 2015.

## Discussion

This paper presents a rule-based alert system for real-time assessment of dengue transmission. It was tailored for Rio de Janeiro, a densely populated city where dengue is highly endemic.

Rio de Janeiro differs from Singapore and other places with ongoing dengue alert systems due to the continuous transmission of dengue (even during winters). From 2010 to Dec 2015, there was not a single week with zero reported cases.

Dengue transmission in Rio de Janeiro is seasonal, modulated by temperature, which affects Aedes aegypti vectorial capacity. Aedes aegypti is found all year round in the city but its abundance varies with temperature (Costa et al, 2015). Honorio et al (2009) found a nonlinear association between temperature and mosquito abundance, with a linear positive association only at temperatures below 22-24°C. Above this temperature, mosquito abundance is high and non sensitive to further increase. This result provides an entomological explanation for the temperature threshold at 22°C found for dengue transmission in the city. In all of these studies, and ours, the strongest association is always with minimum temperature (instead of medium or high temperature). Other meteorological variables, such as humidity and rainfall, are known to affect mosquito biology. Their inclusion in the system is under consideration.

In the literature, there are many proposed early warning systems for dengue. Hii et al (2012) examined the optimal leading time for dengue forecast in Singapore using climate data. They found that a rise of temperature precedes dengue increasing by 1 to 5 months, more strongly with 3-4 months. This approach to modeling dengue, which is commonly used, seeks to associate dengue intensity with temperature, as if they were directly associated. However, biologically speaking, increasing temperature should affect the mosquito abundance and vectorial capacity; in its way, an increased vectorial capacity should affect transmission, that is, the rate of production of new cases. Here, we show that the association between transmission rate and temperature (Rt and temperature) has not such long delay.

Since 2014, outbreaks of an acute exanthematous illness were reported in different parts of the country, mostly diagnosed as dengue. Only in April 2015, Zika virus was detected as the etiological agent. Zika and dengue viruses belong to the flavivirus genus and serological tests do not distinguish between them (Cardoso et al, 2015). In 2015, InfoDengue detected a sustained transmission of dengue starting at April 29 and lasting until June 24. In comparison with previous years, this was a late dengue season, which raised the attention of the city's Dengue Situation Room. Only later, it was confirmed that at least a fraction of these cases were actually Zika infections, what is currently posing a new challenge for the disease notification system.

Desirable features for an online disease alert system are: *Sensitivity*, to detect outbreaks, *speed*, to provide instant information, *stability* to provide comparability with other years and localities, and also *flexibility*, *data quality*, *representativeness*, *acceptability*, *accuracy* and *positive predictive value* (Runge-Ranzinger et al. 2008). The InfoDengue system has provided the city with a faster and more sensitive method for detecting dengue transmission. This is possible because it incorporates climate data which allows detecting favorable transmission conditions before transmission actually starts; and social media data, which allows detection of sudden changes in the social report of dengue symptoms. Still, there is space for further improvement. An investment in better data quality can greatly contributes for the performance of the system. Currently, only four meteorological stations provide temperature data. Satellite data are another potential source of data with better spatial resolution, although not the same temporal resolution. Also, surveillance will gain with a faster notification process, if speed is accompanied by proper digital curation of the data.

Disease alerts are only useful if they trigger actions. For dengue, actions include environmental prophylaxis triggered by an Yellow alert (removal of garbage and covering of containers that can become mosquito breeding sites); mosquito population reduction activities (insecticide, biological control, transgenic mosquitos) triggered by increased transmission (orange alert); and increased medical awareness and health infrastructure for assistance when alert is orange or red. Some of these actions are carried out by health professionals, but the population can also collaborate and demand if the information is made available. Either directly via the site, or indirectly through newspapers (informed by consulting our website), the alert information reached the population.

The adaptation of the Alerta Dengue system to other cities requires a validation of the current set of rules. For similar climates, we expect the same rules will suffice. The expansion of the Alerta Dengue to all 93 cities in Rio de Janeiro's state is mostly complete, exposing some new challenges, for example, the availability and quality of the various data streams, particularly in small communities. In order to accommodate for that we are planning to aggregate multiple small communities into a larger area until it reaches the desired statistical stability. Another source of information, which could be included in the future, is virological surveillance data. We are already working towards integrating entomological surveillance by working with cities that want to start their own system of vector surveillance by means of inexpensive egg traps. Of great importance is the support of public health authorities and their willingness to integrate the results of the Alerta Dengue in their decision making routine. The importance of involvement of local health authorities cannot be overstated, since the maintenance of a fast cycle between data collection and the availability of analytical results, is paramount for the relevance of Alerta Dengue. Also, as we have learned from experience, the definition of a set of well defined alert levels can help turn dengue control more efficient and effective.

Finally, we believe that all the effort invested in combining, cleaning and enriching the various data-streams which feed the Alerta Dengue system, could be of great value as a publicly accessible data source for scholar and health professionals alike. Having more eyes continuously looking at the data can only benefit society's fight to control Dengue and other *Aedes aegypti* borne infections in the long run.

### Support

*Fundação Nacional de Saúde*, FGV, CNPq (CTC, CMD, MG), FAPERJ (CNE CTC)

## Appendix 1. Model for correcting the case counts

Let *Y*_*t*_ be the number of cases occurring at day *t*. At day *τ*>*t*, only a fraction *y*_*t*_(*τ*) of *Y*_*t*_ is known while *x*_*t*_(*τ*) = *Y*_*t*_ – *y*_*t*_(*τ*) is still unknown (censured). From historical data, we have access to uncensored data where we know exactly the time taken for each record to be typed. This dataset is used to compute 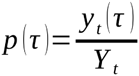 which is the average proportion of cases known as a function of time *τ*. Once this proportion is defined, it can be used to estimate the unknown cases by the following probabilistic model:

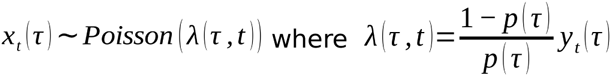

Candidate functions for *p*(*τ*) were the accumulated lognormal, accumulated weibull, logistic and log functions. All functions were fitted to the empirical proportion of cases already notified at delay *τ* using the survival library in R (R Core Team, 2015; Therneau, 2015) and the best model (lognormal) chosen by AIC. The fitted function was *p*(*τ*)=*Φ*(*τ*, *mean* = 3, *var* = 0.91) where Φ is the lognormal function.

**Figure S1.**
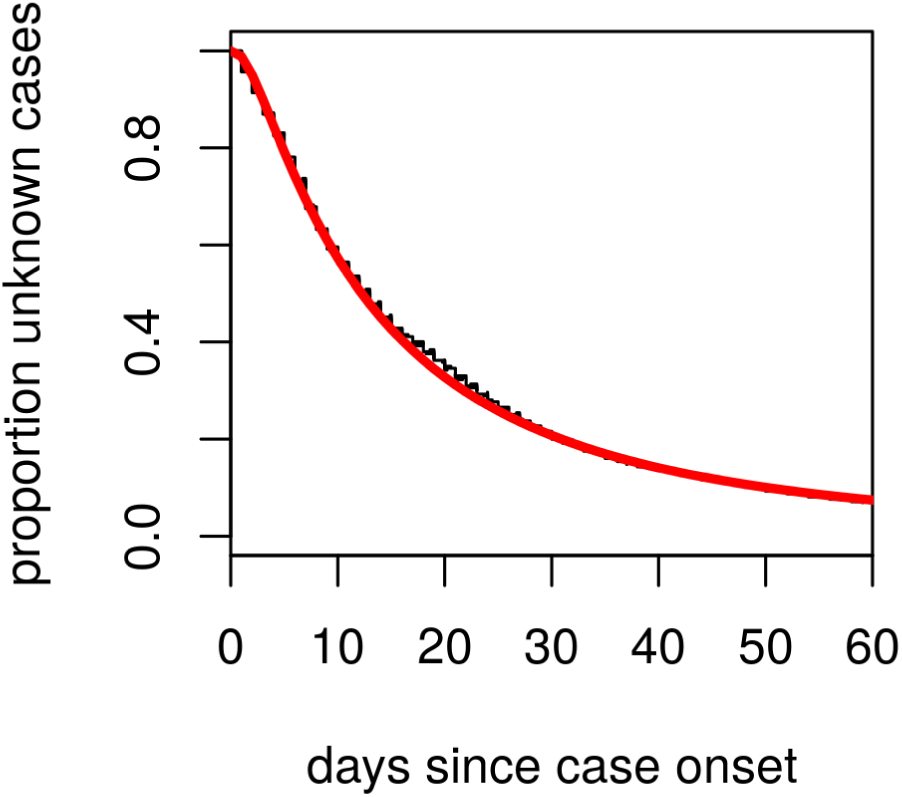
Black: observed proportion of cases still not typed days below the onset. Red: fitted lognormal function.

To test the procedure, we created an artificial time series containing only records that where known within two weeks from occurrence. Using the notation of the model, this corresponds to *y*_*t*_(*τ* = 2). The solid black line is *Y*_*t*_, the total cases that we want to predict. The predicted number of cases (in green) shows good agreement with the observed cases, suggesting that this approach is adequate for case estimation.

